# DNA methylation in the APOE gene: its link with Alzheimer’s and cardiovascular health

**DOI:** 10.1101/811224

**Authors:** Jure Mur, Daniel L. McCartney, Rosie M. Walker, Archie Campbell, Mairead L. Bermingham, Stewart W. Morris, David J. Porteous, Andrew M. McIntosh, Ian J. Deary, Kathryn L. Evans, Riccardo E. Marioni

## Abstract

Genetic variation in the apolipoprotein E (*APOE*) gene is associated with Alzheimer’s disease (AD) and risk factors for cardiovascular disease (CVD). DNA methylation at *APOE* has been linked to altered cognition and AD. It is unclear if epigenetic marks could be used for predicting future disease. We assessed blood-based DNA methylation at 13 CpGs in the *APOE* gene in 5828 participants from the Generation Scotland (GS) cohort. Using linear regression, we examined the relationship between *APOE* methylation, cognition, cholesterol, and the risks for AD and CVD. DNA methylation at two CpGs was associated with the ratio of total-to-HDL cholesterol, but not with cognition, or the risks of AD or CVD. *APOE* methylation could be involved in the levels of blood cholesterol, but there is no evidence for the utility of *APOE* methylation as a biomarker for predicting AD or CVD.

## 1. Introduction

### 1.1. The APOE gene

The apolipoprotein E (*APOE)* gene is located on chromosome 19, with two SNPs in its fourth exon defining three alleles, ε2, ε3, and ε4, resulting in the production of three isoforms of the apoE protein, apoE2, E3, and E4 [1]. The *APOE* ε4 genotype is a well-known risk factor for Alzheimer’s disease (AD) [2,3] and a prominent candidate in cardiovascular research [4,5]. The protein is expressed in various tissues, including the brain [6]. It acts as a ligand for members of the low-density lipoprotein (LDL) receptor family and is involved in the clearance of lipoproteins and cholesterol from the circulatory system [7,8].

*APOE* exhibits an allele-specific association with risk of AD. Possession of the ε4-allele confers an increased risk, while the ε2-allele is protective [2]. The isoforms of apoE differ in their binding-affinity for lipoproteins and LDL-receptors, and differentially influence levels of serum cholesterol [9]. The Ε2-variant reduces the levels of total- and LDL-cholesterol, while the E4 variant raises them [6]. However, the connection between *APOE* and cardiovascular disease (CVD) is less robust. While there is some evidence for an increased risk of CVD-related death in carriers of the ε4-allele [1], this has not always been observed [10].

### 1.2. APOE methylation in AD and CVD

Epigenetic modifications have been associated with many human disorders [11]. DNA methylation is the addition of a methyl group to the 5-position of cytosine in the context of a cytosine-guanine dinucleotide (CpG; [12]). Many studies have found a link between AD and methylation [12,13]. Candidate gene studies for *APOE* have shown that methylation in this gene is associated with dementia [14,15,16,17] with neuritic amyloid plaque burden [18], and with cognitive ability [19].

Associations have been suggested between epigenetic mechanisms and risk factors for cardiovascular disease [20,21,22,23]. Whereas a recent systematic review identified 34 candidate-gene studies on methylation in CVD [4], few have explored modifications in *APOE*, and there have been some conflicting results: while Karlsson et al. [15] found no evidence for differences in *APOE* methylation in blood between patients with CVD and healthy controls, Ji et al. [24] reported *APOE* hypermethylation in the blood of patients with coronary heart disease compared to controls.

To help clarify this role of *APOE*, we utilize data from Generation Scotland, a large population-based cohort, the size of which is several times greater than samples in previous studies, providing a robust analysis of *APOE* methylation and cognitive and vascular health. We characterise blood-based methylation in the promoter region, 2^nd^ and 3^rd^ exons and introns, and 4^th^ exon of the *APOE* gene, and explore links between *APOE* methylation and a variety of markers of cognitive function, AD, vascular health, and CVD. Due to the importance of developing new clinical biomarkers for health outcomes before the onset of disease, we restrict our sample to ages between 30 and 65 years.

## 2. Methods

### 2.1. The sample

Generation Scotland (GS; [25,26]) is a family-based cohort of over 22,000 individuals from Scotland (aged 18-99 years) that were genotyped and extensively phenotyped during the baseline assessment between the years 2006 and 2011. Blood-based genome-wide DNA-methylation were generated in 9551 individuals as part of the sub-study Stratifying Resilience and Depression Longitudinally (STRADL; [27]).

### 2.2. Methylation and APOE measurements

DNA methylation in peripheral blood samples was profiled in two sets (Set 1: n = 5,087 in 2017 and Set 2: n = 4,450 in 2019), using the Illumina HumanMethylationEPIC BeadChip as described previously [28]. Briefly, low quality samples, probes with low detection rates, and participants for which the predicted sex did not match the recorded sex were excluded (**Supp. Methods 1**).

We removed related participants from Set 1 using a genetically determined relationship cut-off of >0.05 (GCTA GREML; [29]) to reduce the potential influence of shared genetics on the findings. The participants in Set 2 were unrelated to each other and to those in Set 1. We restricted the analysis to CpGs located on chromosome 19 between 45,409,039 - 45,412,650bp, which corresponded to the region encompassing the *APOE*-gene (UCSC GRCh37/hg19 genome build). To restrict the age of our sample, we removed participants younger than 30 years and older than 65 years. After combining the two sets, our final sample consisted of a total of 13 CpGs in 5,828 participants (**Supp. Figure 1**).

*APOE*-haplotype status was determined using Taqman technology at the Clinical Research Facility, Western General Hospital, Edinburgh. Based on the identity of the nucleotides at SNP-positions rs429358 and rs7412, participants with the ε3/ε3-haplotype were classified as ε3-carriers, participants with the ε2/ε2- and ε2/ε3-haplotypes were classified as ε2-carriers, and participants with the ε3/ε4- and ε4/ε4-haplotypes were classified as ε4-carriers. The 126 participants (2.2%) with the ε2/ε4-genotype were not included in analyses in which carrier-status was implemented as a variable.

### 2.3. Characterisation of APOE methylation

We examined the methylation status of 13 CpGs in the *APOE* gene (**Figure 1A**). Twelve CpGs exhibited similar methylation levels to those previously described (**Supp. Table 1**; [15,16,22]), whereas one of them (cg20051876, which is unique to the Illumine EPIC array) had not been reported before. Based on the Illumina-annotated locations of the CpGs on the chromosome, their relative distances to each other, and their methylation levels, the CpGs were classified into three groups: hypermethylated (each site > 50% mean methylation) and lying in the promoter region (region 1: cg20051876, cg14123992, cg04406254), hypomethylated (each site < 50% mean methylation) and lying in the region encompassing the first two exons and introns (region 2: cg26190885, cg12049787, cg08955609, cg18768621, cg19514613, cg06750524), and hypermethylated and lying in the 4^th^ exon (region 3: cg16471933, cg05501958, cg18799241, cg21879725).

**Figure 1:**
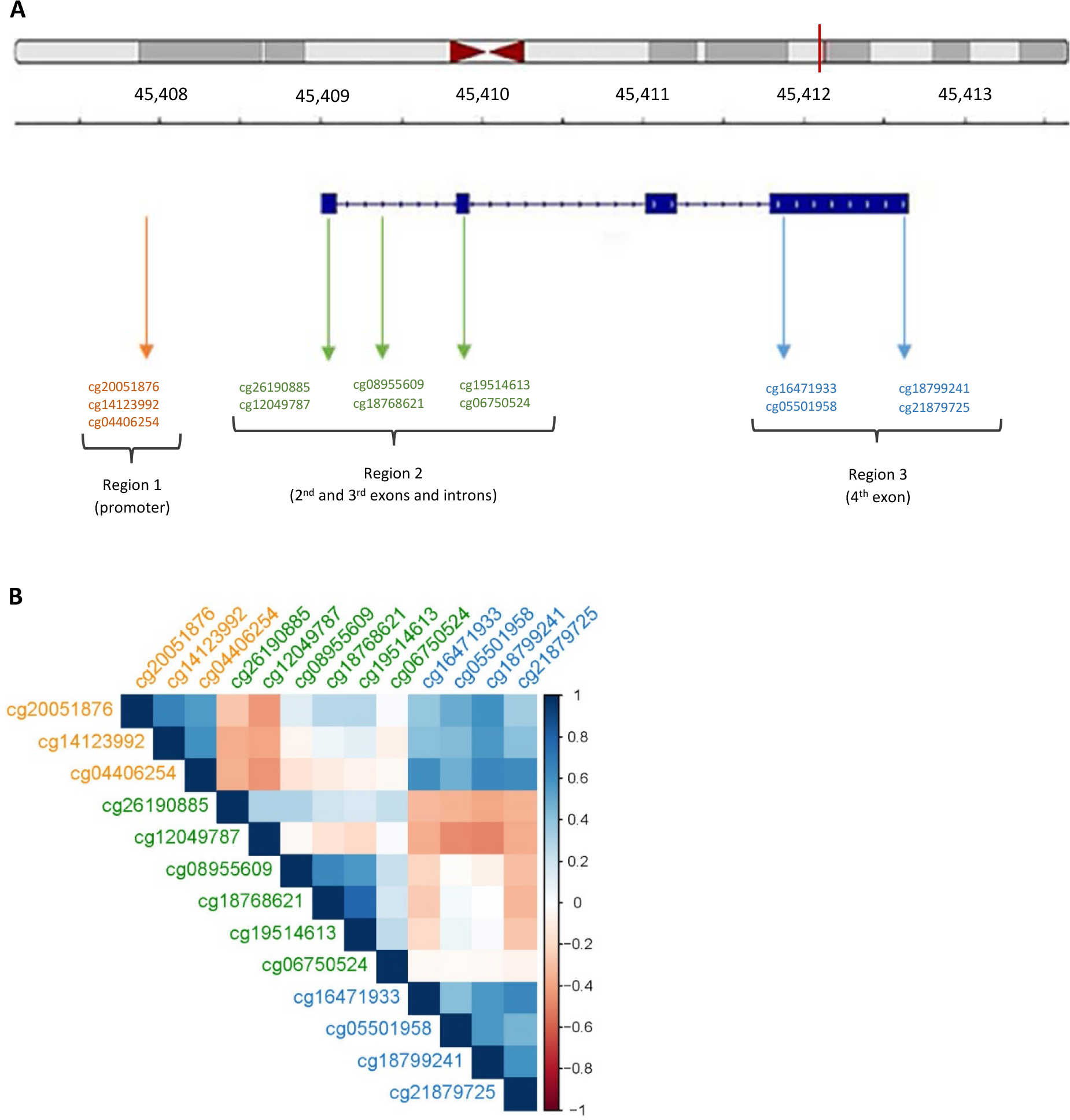
Structure of the *APOE* locus and the location of CpGs (**A**), and correlations among CpGs in the *APOE* gene (**B**). **A**: The top panel shows the chromosomal location of the *APOE* gene; the second panel represents the location on the chromosome in kb; the third panel shows the structure of the *APOE* gene, with exon-regions thickened. The approximate locations of CpGs on the *APOE* gene is indicated by the arrows, with colours representing regional classification: region 1 (orange), region 2 (green), or region 3 (blue). **B**: The strength of the correlation is represented by a colour code, ranging from dark red (strong negative correlation) to dark blue (strong positive correlation). The colour of the text denotes the location for a given CpG on the *APOE* gene: region 1 (orange), region 2 (green), or region 3 (blue).

### 2.4. Cognitive variables

Four measures of cognitive function [26] were evaluated: (1) the Wechsler digit symbol test as a measure of the speed of information processing, (2) the letter-based (C, F, L) phonemic verbal fluency test as a measure of executive function, (3) the Mill Hill Vocabulary test (combined junior and senior synonyms) as a measure of crystallised ability, and (4) the Wechsler immediate and delayed Logical Memory tests (one story) as a measure of verbal declarative memory. Participants with a zero-value were judged as resulting from non-participation and were recorded as missing. A measure of global cognitive function (cognitive-g) was derived by applying principal component analysis (PCA) to the four cognitive measures, with cognitive-g defined as the first unrotated principal component. This factor explained 44% of the total variance from all four cognitive measures, with loading values ranging between 0.47 and 0.53.

### 2.5. Approximations for the risk of AD and of CVD

Only three participants reported having AD, and 203 participants (3.5%) reported having CVD. We therefore created approximate measures of risk that were based on reports by participants about these illnesses for close relatives (parents, siblings, and grandparents). For both AD and CVD, each relative was assigned either 0 or 1 (0: absence of the disorder, 1: presence of the disorder) based on the report of the participant. To derive each risk measure, we calculated the weighted sums of family records for each participant as follows:

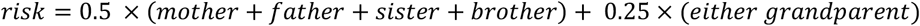

The choice of weights used for a given family member was based on kinship/relatedness between that relative and the participant. The participants that self-reported as having AD or CVD were excluded from the models that included the risk of AD or the risk of CVD as predictor variables, respectively. Due to a low number of relatives with AD and a consequential skew in the distribution of AD-risk, AD-risk was transformed into a categorical variable AD-class (low risk: no close relatives with AD, n = 4854; high risk: at least one close relative with AD, n = 974). For cholesterol levels, the total-to-HDL ratio was used.

### 2.6. Covariates

For alcohol consumption, we did not include participants that have quit drinking (n = 373) or whose answers relating to recent alcohol consumption according to their own assessment did not correspond to their usual drinking pattern (n = 1557). Information on smoking was processed as described previously [30]; we did not include participants that have quit smoking (n = 673). The sample sizes for the fully-adjusted models (see below) thus depended on the included covariates with missing values. Smoking was assessed in pack years and alcohol consumption in units per week. Activity levels were recorded in minutes per week; two different versions of the questionnaire were used in the study, with each participant completing only one of them. We therefore combined the two versions so that the responses by all participants were on the same scale (**Supp. Methods 2**).

### 2.7. Statistical analysis

For all numerical variables, outliers – defined as scores beyond four interquartile ranges from the median – were excluded. For each analysis, linear mixed models were built. For each CpG (outcome), two models were run: (1) a basic-adjusted model containing the primary predictor variables as defined in the hypothesis and (2) a fully-adjusted model that included additional covariates:

Basic-adjusted model:

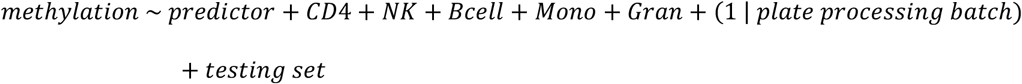
Fully-adjusted model:

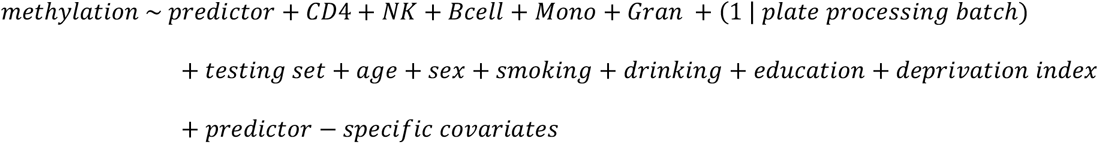

The predictor-specific covariates used in the fully-adjusted model varied according to the predictor variable of the model and, whenever possible, corresponded with previous studies [15,22]. Both the basic- and the fully-adjusted model included as covariates the batch in which the samples of a given participant were processed on the array (fitted as a random effect on the intercept) and the estimated proportions of CD8+− and CD4+− T-cells, natural killer cells, B cells, monocytes, and granulocytes. Cell composition correction controls for the fact that DNA methylation patterns can be confounded by the heterogeneity of cells in the tissue used for analysis [31].

The threshold for statistical significance was Bonferroni-corrected for each CpG. All continuous variables (except the variable *age* when used as a predictor) were transformed to have a mean of 0 and a standard deviation of 1. When the predictor was a categorical variable, analysis of variance was performed, comparing the model of interest with the same model excluding the predictor variable.

All statistical analyses were performed in R.

## 3. Results

### 3.1. Sample characteristics

Among the 5828 participants, 3399 (58.3%) were female and 2429 (41.7%) were male. The age range was 30-65 years and the mean age was 52.7 years (**Table 1**). *APOE* genotype frequencies were comparable to those described previously for the British population (**Table 1**; [10,32]). Correlations between DNA methylation levels in blood (this study) and brain tissue (publicly available datasets) for the 13 CpGs in this study ranged from −0.30 to 0.51 for the whole brain (IMAGE-CpG: https://han-lab.org/methylation/default/imageCpG, [33], last accessed 13.10.2019), from −0.26 to 0.40 for Brodmann area 20 (BECon: https://redgar598.shinyapps.io/BECon/, [34], last accessed 13.10.2019), and from −0.21 to 0.20 for the entorhinal cortex (https://epigenetics.essex.ac.uk/bloodbrain/, [35], last accessed 13.10.2019), with most CpGs exhibiting low correlations between brain- and blood methylation (**Supp. Table 1**).

**Table 1:** Descriptive statistics and frequencies of genotypes and alleles/carriers in the sample.

| Trait | Mean<sup>*</sup> | SD<sup>†</sup> |
| --- | --- | --- |
| Age | 53.0* | 14.0* |
| Years in education<sup>‡</sup> | 4.0* | 3.0* |
| Deprivation quintile<sup>§</sup> | 4.0* | 3.0* |
| Processing speed (Wechsler digit symbol test) | 71.2 | 15.6 |
| Executive function (letter-based verbal fluency test) | 41.1 | 11.7 |
| Verbal ability (Mill Hill Vocabulary test) | 30.1 | 4.5 |
| Verbal declarative memory (Wechsler logical memory test immediate and delayed) | 30.9 | 7.7 |
| Heart rate (beats/minute) | 63.7 | 10.3 |
| Ratio of total-to-HDL cholesterol | 3.7* | 1.6* |
| Alcohol units per week | 8.0* | 12.0* |

| Trait | N | % |
| --- | --- | --- |
| Sex (female) | 3399 | 58.3 |
| AD in close relatives | 974 | 16.7 |
| CVD in close relatives | 3474 | 59.6 |

| Genotype/carrier | n | % |
| --- | --- | --- |
| ε2/ ε2 | 31 | 0.5 |
| ε2/ ε3 | 647 | 11.4 |
| ε2/ ε4 | 126 | 2.2 |
| ε3/ ε3 | 3352 | 59.0 |
| ε3/ ε4 | 1370 | 24.1 |
| ε4/ ε4 | 156 | 2.7 |
| ε2-carrier | 678 | 12.2 |
| ε3-carrier | 3352 | 60.3 |
| ε4-carrier | 1526 | 27.5 |

\* The median is given when appropriate (denoted with \*).
† The interquartile range is given when appropriate (denoted with †).
‡ A 10-grade classification-system, where 0: 0 years, 1: 1-4 years, 2: 5-9 years, 3: 10-11 years, 4: 12-13 years, 5: 14-15 years, 6: 16-17 years, 7: 18-19 years, 8: 20-21 years, 9: 22-23 years, 10: ≥ 24 years.
§ Scottish Index of Multiple Deprivation (SIMD), 2011.

### 3.2. Association between CpGs

The correlations in methylation for all pairwise combinations between the 13 CpGs ranged from −0.49 to 0.78, with a mean absolute correlation of 0.32 (**Figure 1B**). After correcting for multiple comparisons, 68/78 correlations were statistically significant at p<6.4×10^−4^ (=0.05/78), with 38 (60%) positive, and 30 (40%) negative. CpGs within a region tended to be methylated to a similar extent: all significant correlations among CpGs within region 1 (3/3, 100%) and within region 3 (6/6, 100%), and most of the significant correlations among CpGs within region 2 (11/13, 85%) were positive. Moreover, there was a tendency for CpGs in regions 2 and 3 to be methylated in opposite directions: most of the significant correlations (17/22, 77%) were negative. In contrast, regions 1 and 3 tended to be methylated in the same direction, with all significant correlations (12/12, 100%) positive.

### 3.3. Association between methylation and genotype

After correcting for multiple testing, there was evidence for an association between *APOE* carrier-status and DNA methylation at five CpGs in the fully-adjusted models: cg14123992 (η^2^=0.003, p=3.9×10^−4^), cg04406254 (η^2^=0.004 p=4.0×10^−6^), cg06750524 (η^2^=0.022, p=8.6×10^−16^), cg16471933 (η^2^= 0.014, p=2.4×10^−16^), and cg21879725 (η^2^=0.003, p=1.3×10^−4^) (**Figure 2**, **Supp. Table 2**). Post-hoc testing between the *APOE* groups showed that at all five CpGs the associations were due to higher methylation levels in ε4 carriers compared to ε3 carriers.

**Figure 2:**
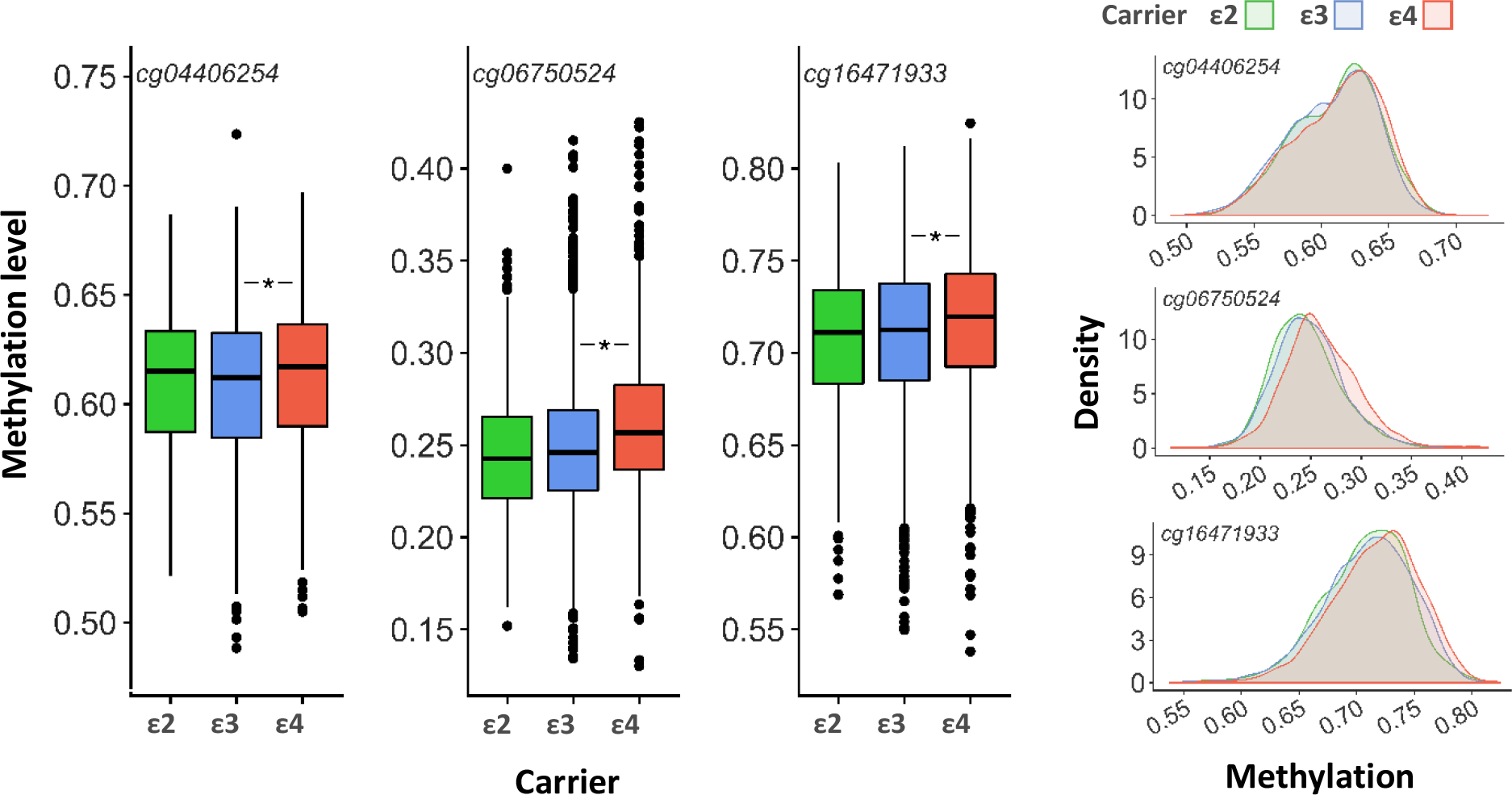
Boxplots (**left**) and density plots (**right**) for differences in methylation levels between groups with different carrier-status. The statistically significant differences between groups after adjusting for multiple comparisons (p < 0.0035) are denoted with asterisks. Depicted are only CpGs for which the fully-adjusted mixed linear models and post-hoc tests between carriers were significant after adjusting for multiple testing.

### 3.4. Age-dependent drift in DNA-methylation

In the basic-adjusted model, 7/13 CpGs showed an association with age after correction for multiple testing (R^2^ range: 4.0×10^−5^ – 0.03; **Supp Table 3**). Three CpGs remained significant after correction for multiple testing in the fully-adjusted model (**Figure 3**, **Supp. Table 3**). Most CpGs within a region exhibited the same direction of change with respect to aging. CpGs in the generally hypomethylated region 2 tended to have greater methylation with older age, whereas CpGs in the generally hypermethylated regions 1 and 3 tended to have lower methylation with older age. None of the CpGs in our analyses were in a list of age-related variable methylated positions (aVMPs; [36]). However, the Breusch-Pagan test for heteroscedasticity indicated non-random variation in the residuals by age for all CpGs in both the basic- and the fully-adjusted models (**Supp. Table 4**).

**Figure 3:**
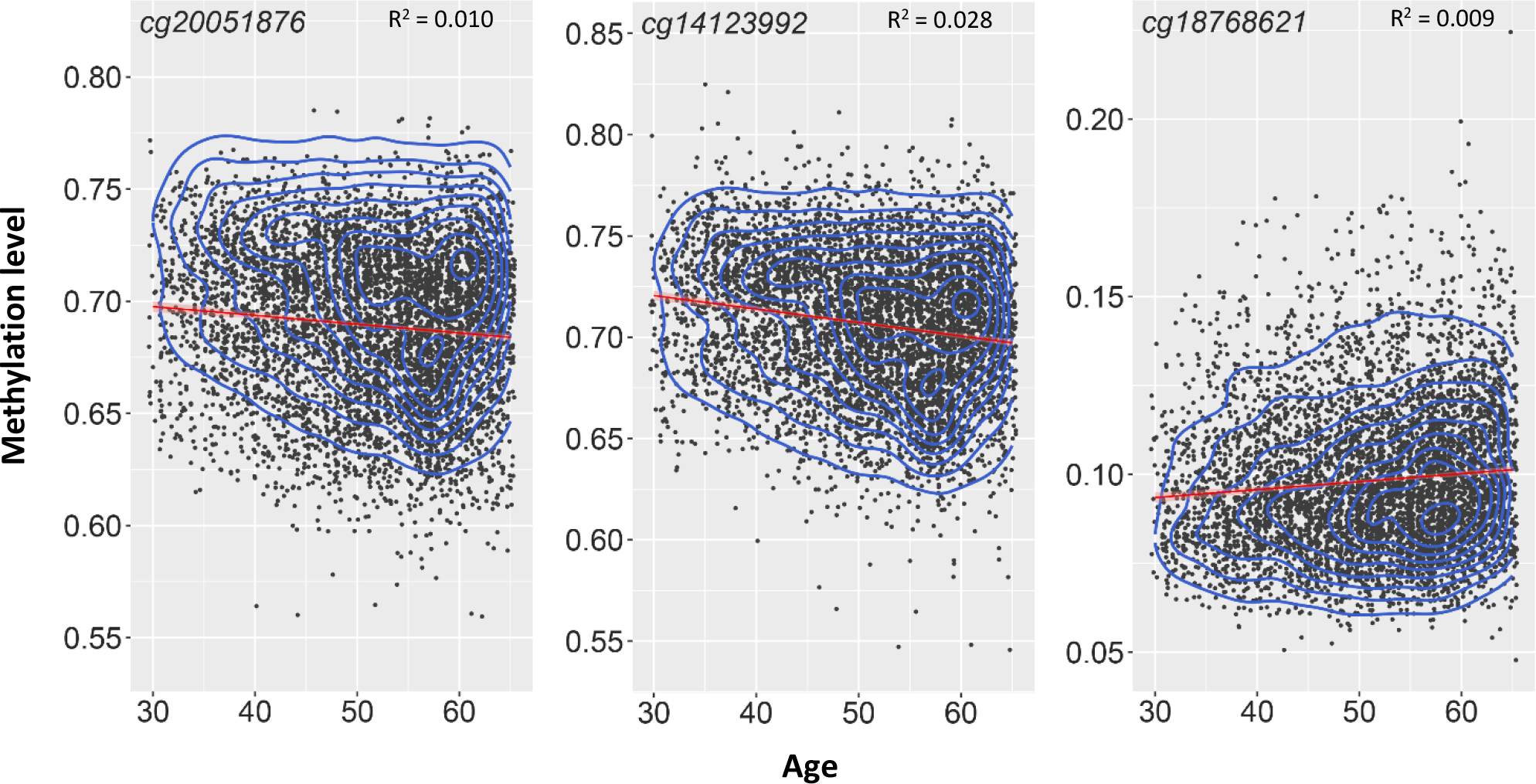
Relationship between age and methylation; the red line represents the linear fit between the two variables, the blue contour lines represent the density of the observations. Depicted are CpGs cg20051876, cg14123992, and cg18768621, for which the fully-adjusted mixed linear model was significant.

### 3.5. APOE methylation and cognitive function

We observed no association between general cognitive ability and DNA methylation in the fully-adjusted regression models after correction for multiple testing (**Supp. Table 5**). There were associations between the individual cognitive tests and DNA methylation levels in the basic- but not fully-adjusted regression models (**Supp. Table 6**).

### 3.6. APOE methylation and the risk of AD

To determine whether carrier-status was associated with AD-class (0: absence of the disorder, 1: presence of the disorder), a logistic regression was run, with AD-class as the outcome variable and carrier-status as the predictor. We observed no associations between DNA methylation levels and AD-risk in the basic- or fully-adjusted models after correcting for multiple testing (**Supp. Table 7**). The Wald Chi-Squared test confirmed a general effect of carrier-status in the basic-adjusted model (*χ*^2^=49.8, p=8.9×10^−11^) and in the fully-adjusted model (*χ*^2^=33.4, p=2.6×10^−7^). The ε4-allele was significantly associated with the high-risk AD-class both in the basic- (OR=1.70, p=3.2×10^−11^) and fully-adjusted models (OR=1.77, p=1.2×10^−7^).

### 3.7. APOE methylation and the risk of CVD

We observed no association between *APOE* methylation and the risk measure of CVD, nor was there an association between *APOE* genotype and the risk measure of CVD (**Supp. Table 8**). However, *APOE* genotype was linked to the total-to-HDL cholesterol ratio in the fully-adjusted model (*χ*^2^=68.5, p=1.3×10^−15^). Specifically, both the ε2- and ε4-carriers differed in the ratio of total-to-HDL cholesterol compared to ε3-carriers in the fully-adjusted model (ε2: estimate=−0.22, SE=0.048, p=6.4×10^−6^; ε4: estimate=0.21, SE=0.036, p=1.2×10^−8^). The ratio of total-to-HDL cholesterol was significantly associated with methylation levels after correcting for multiple testing at three CpGs in the fully-adjusted models: cg08955609, cg18768621, and cg16471933 (**Supp. Table 9**). Due to the strong association between *APOE* genotype and cholesterol levels, the fully-adjusted model was rerun with *APOE*-genotype as covariate; cg08955609 (estimate=−0.065, SE=0.019, p=6.7×10^−4^) and cg18768621 (estimate=−0.059, SE=0.017, p=4.3×10^−4^) remained statistically significant (**Figure 4**, **Supp. Table 9**). The validity of CVD-risk as a risk measure for CVD was confirmed by running a linear regression model, with self-reported CVD as an outcome measure and CVD-risk as a predictor variable, with covariates as in the fully-adjusted model above (estimate=0.02, error=0.0032, p=2.1×10^−11^).

**Figure 4:**
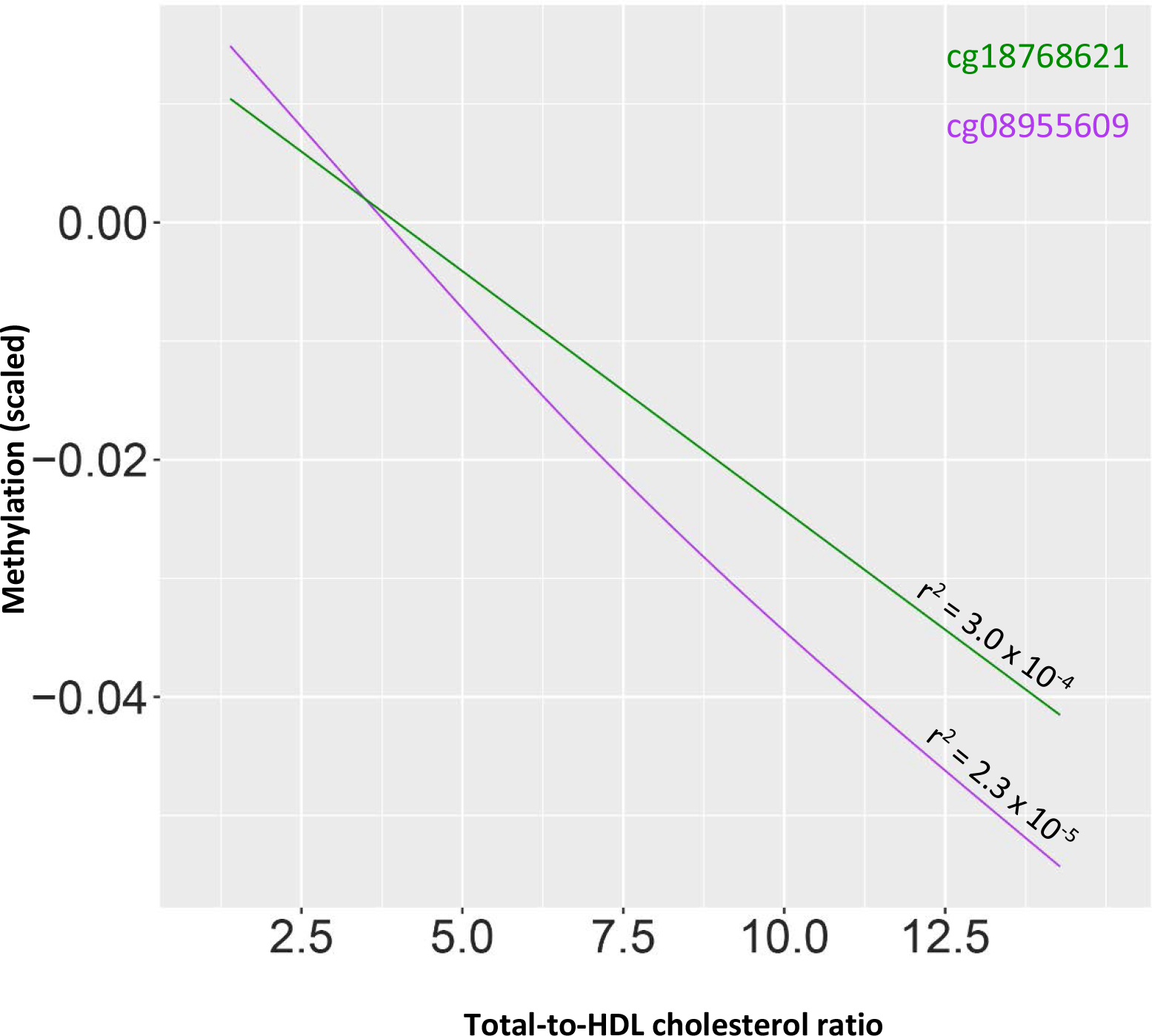
Relationship between total-to-HDL cholesterol ratio and methylation. Depicted are CpGs cg08955609 and cg18768621, for which the linear models were significant.

## 4. Discussion

### 4.1. Correlations between CpGs and age-drift in methylation

In this study, we used DNA methylation- and phenotypic data from a large cohort, Generation Scotland, to explore methylation in the *APOE* gene and its association with risk factors for AD, CVD, and blood cholesterol.

We observed correlations among CpGs which had been reported before [19]. Compared with Liu et al. [19], the correlations observed in our study were stronger and more of them were negative. In contrast to the findings of Karlsson et al. [15], the methylation levels of five CpGs correlated with carrier-status for the different *APOE* alleles. This difference may be due to the increased power of our study over that of Karlsson et al. [15].

We replicated a finding by Ma et al. [22] of age-drift in methylation in the *APOE* gene: 3/13 CpGs showed age-dependent changes. In each hypermethylated CpG, methylation levels tended to decrease with age, while in each hypomethylated CpG, methylation levels tended to increase with age. All CpGs exhibited heteroscedasticity for the change in methylation as a function of age, demonstrating that the drift could be due to increases in methylomic variability with increasing age.

### 4.2. Relationship between APOE methylation, AD and cognition

Neither cognition, nor family history of AD were associated with *APOE* methylation. This might be due to a lack of appreciable changes in *APOE* methylation before the onset of symptoms of AD. Most previous studies that linked differential *APOE* methylation with AD were conducted on tissue from patients that had either been screened positive for cognitive dysfunction [15], or diagnosed with AD [14,16,17]; only one study [19] evaluated the relationship between DNA methylation and cognition in healthy participants. The epigenetic changes in *APOE* accompanying AD-related cognitive decline could result from the pathophysiology of the disorder or from adaptive responses of the organism as a result of AD, neither of which may be present at appreciable levels in the population studied here. Another consideration concerns the tissue used. Foraker et al. [14] used post-mortem brain-tissue, Karlsson et al. [15] and Liu et al. [19] used blood, while Shao et al. [16] and Wang et al. [17] used both. In fact, the latter were not able to replicate their findings from brain-tissue in blood. Blood represents an attractive medium for identifying biomarkers for disease. Indeed, it has been reported that patients with AD and healthy controls can be distinguished based on gene-expression patterns in blood [37]. However, the *APOE* gene is differently expressed between brain tissue and blood [16,17] and *APOE* CpGs exhibit relatively modest correlations between blood- and brain methylation [33,34,35]. Thus, blood-methylation may exhibit AD-associated changes that are distinct from methylation changes in the brain or they might appear later in the course of the disorder. Finally, while some studies have reported AD-associated changes in *APOE* methylation as described above, little research has been done on the topic and few – if any – replication studies have been performed to validate the effects. Moreover, some prominent studies that investigated associations between DNA methylation and AD across the entire genome did not report *APOE* to be altered in the disorder [13,38].

### 4.3. Relationship between APOE methylation and blood cholesterol

We did not replicate the finding [24] of an association between *APOE* methylation and CVD; our results are in line with reports by Karlsson et al. [15] and Sharma et al. [39]. However, we did find a negative association between *APOE* methylation and the ratio of the total to HDL-cholesterol at cg08955609 and cg18768621; a finding that – to our knowledge – had not been reported before. Our measurements of cholesterol provide only levels of total blood cholesterol and of HDL cholesterol. Because of this – especially considering the opposite correlation of *APOE* genotype with the concentrations of LDL and VLDL, respectively [40] – it is not possible to assess the relationship between methylation and the individual components of blood cholesterol.

### 4.4. Limitations and future directions

The results of the present study offer additional insight into the epigenetics of the *APOE* gene and its association with relevant phenotypes. The main strength of the study is the large sample size from a relatively representative sample of the Scottish population. Nevertheless, we recognize several limitations. Firstly, the measures of risk for AD and CVD were inferred from information on participants’ relatives. The rough approximation of risk underestimates the importance of environmental factors. Moreover, different participants in our cohort have different numbers of relatives. Secondly, the study is limited to (a subset of) the methylome; other important epigenetic mechanisms that are currently less accessible to analysis at scale might also play important roles in the studied interactions. Thirdly, our study is based on methylation data from the blood, which for most CpGs correspond to methylation patterns in the brain. This might preclude the identification of potential epigenetic changes prior to disease onset and allows only limited insight into underlying biological processes. However, due to the multitude of peripheral biological processes associated with AD, blood may be a legitimate tissue for DNA methylation studies. Finally, this study adopts a candidate gene approach; while *APOE* is a biologically plausible candidate for implications in AD and CVD, genome-wide approaches may inform on the relative importance of *APOE*.

In conclusion, we showed that CpGs in the *APOE* gene exhibit correlations within- and between distinct regions of the gene and that the methylation levels at some CpGs of the *APOE* gene correlate with the *APOE* genotype. Furthermore, we found an association between methylation level at cg8955609 and cg18768621, and blood cholesterol. We did not find differences in the levels of *APOE* methylation between individuals at low risk and individuals at high risk of developing either AD or CVD. Future work might explore longitudinal changes in methylation as it relates to adverse health outcomes to circumvent the problems of the present study and improve our understanding of the exact timing of epigenetic changes in AD and CVD.

## Acknowledgements

GS received core support from the Chief Scientist Office of the Scottish Government Health Directorates (CZD/16/6) and the Scottish Funding Council (HR03006). Genotyping and DNA methylation profiling of the GS samples was carried out by the Genetics Core Laboratory at the Clinical Research Facility, University of Edinburgh, Edinburgh, Scotland and was funded by the Medical Research Council UK and the Wellcome Trust (Wellcome Trust Strategic Award “STratifying Resilience and Depression Longitudinally” ([STRADL; Reference 104036/Z/14/Z]). DLM and REM are supported by Alzheimer’s Research UK major project grant ARUK-PG2017B-10. IJD is a member of the Lothian Birth Cohorts group at the University of Edinburgh and is supported by Age UK (Disconnected Mind grant), the Medical Research Council (MR/R024065/1), and the USA’s National Institutes of Health (1RO1AG054628-01A1). JM is supported by funding from the Wellcome Trust 4-year PhD in Translational Neuroscience – training the next generation of basic neuroscientists to embrace clinical research [108890/Z/15/Z].

