## Supplementary material for "DNA methylation in the APOE gene: its link with Alzheimer’s and cardiovascular health": Supp. material

**Supp. Methods 1**: *Quality control steps for DNA methylation.*
After plotting the log median intensity of methylated vs. unmethylated signal per array, outliers were excluded by visual inspection. Samples with ≥ 1% of cytosine-guanine dinucleotides with a detection p value greater than 0.05, probes with a bead-count of less than 3 in more than 5 samples, and probes with ≥ 0.5% of samples with a detection value greater than 0.05 were excluded. Additionally, samples in which predicted sex did not match recorded sex were excluded.

**Supp. Methods 2**: *Combination of activity data.*
The two questionnaires on activity (v2 and v5) differed in the following ways: v2 differentiated between work-related and non-work-related physical activity, while v5 did not. v2 had three different categories of physical activity (*very active, moderately active, inactive*), while v5 had four (*vigorous activity, moderate activity, walking, sitting*). In v2, we first combined work- and non-work-related activity; in v5, we combined *walking* with *moderate activity*, so that both versions of the questionnaire had analogous coding of the data. Moreover, all physical activity was re-calculated into minutes/week. To generate a single metric for physical activity, PCA was applied to the three categories of physical activity, with general physical activity defined as the first unrotated principal component.

**Supp. Figure 1**: Flow chart of the DNA methylation analysis pipeline.

Remove
irrelevant CpGs

Remove SNP-probes

Remove
related participants

Remove participants outside the age
range

**Supp. Table 1**: Average methylation levels, standard deviations in methylation levels, locations of CpGs on the chromosome, rho-values for the correlation in methylation between blood and whole brain, between blood and Brodmann area 20, and between blood and the entorhinal cortex. The CpGs are classified into region 1 (orange), region 2 (green), or region 3 (blue).

| **CpG** | **Mean** | **SD** | **Location on chromosome (kb)** | **Blood-brain association** | | |
| --- | --- | --- | --- | --- | --- | --- |
|  |  |  |  | **Whole brain [33]** | **BA20  [34]** | **Entorhinal cortex [35]** |
| cg20051876 | 0.69 | 0.035 | 45,407,860 | n.a. | n.a. | n.a. |
| cg14123992 | 0.71 | 0.036 | 45,407,868 | 0.18 | -0.26 | -0.15 |
| cg04406254 | 0.61 | 0.032 | 45,407,945 | 0.45 | 0.11 | 0.16 |
| cg26190885 | 0.09 | 0.014 | 45,409,005 | -0.1 | 0.14 | 0.05 |
| cg12049787 | 0.08 | 0.018 | 45,409,080 | 0.18 | 0.29 | 0.13 |
| cg08955609 | 0.05 | 0.009 | 45,409,353 | 0.51 | 0.12 | 0.20 |
| cg18768621 | 0.10 | 0.021 | 45,409,440 | -0.2 | -0.03 | n.a. |
| cg19514613 | 0.13 | 0.024 | 45,409,713 | -0.3 | -0.18 | -0.12 |
| cg06750524 | 0.25 | 0.036 | 45,409,955 | 0.0 | 0.21 | 0.14 |
| cg16471933 | 0.71 | 0.038 | 45,411,802 | 0.24 | -0.26 | -0.12 |
| cg05501958 | 0.94 | 0.014 | 45,411,873 | 0.04 | -0.08 | -0.05 |
| cg18799241 | 0.80 | 0.031 | 45,412,599 | n.a. | 0.08 | 0.003 |
| cg21879725 | 0.74 | 0.033 | 45,412,647 | -0.3 | -0.05 | -0.21 |

**Supp. Table 2:** Basic- and adjusted linear mixed models with methylation level as the outcome variable and carrier-status as the predictor variable. Bold font denotes significant values after correcting for multiple testing (p < 0.0038). The CpGs are classified into region 1 (orange), region 2 (green), or region 3 (blue).

|  | **Basic-adjusted model** | | | **Fully-adjusted model** | |  |
| --- | --- | --- | --- | --- | --- | --- |
| **CpG** | ***χ*^2^** | **p** | **η^2^** | ***χ*^2^** | **p** | **η^2^** |
| cg20051876 | 6.2 | 0.045 | 0.001 | 4.5 | 0.11 | 0.001 |
| **cg14123992** | **14.9** | **5.6 x 10^-4^** | **0.001** | **15.7** | **3.9 x 10^-4^** | **0.003** |
| **cg04406254** | **25.4** | **3.1 x 10^-6^** | **0.002** | **24.9** | **4.0 x 10^-6^** | **0.004** |
| cg26190885 | 0.6 | 0.73 | 0.000 | 0.01 | 1.0 | 0.000 |
| cg12049787 | 0.6 | 0.74 | 0.000 | 0.5 | 0.78 | 0.000 |
| cg08955609 | 0.3 | 0.86 | 0.000 | 0.04 | 0.98 | 0.000 |
| cg18768621 | 0.2 | 0.89 | 0.000 | 0.2 | 0.90 | 0.000 |
| cg19514613 | 10.5 | 0.005 | 0.002 | 5.6 | 0.06 | 0.002 |
| **cg06750524** | **144.0** | **<2.2 x 10^-16^** | **0.025** | **69.4** | **8.6 x 10^-16^** | **0.022** |
| **cg16471933** | **81.4** | **<2.2 x 10^-16^** | **0.009** | **71.9** | **2.4 x 10^-16^** | **0.014** |
| cg05501958 | 0.8 | 0.67 | 0.000 | 1.2 | 0.54 | 0.000 |
| cg18799241 | 5.6 | 0.06 | 0.000 | 5.7 | 0.06 | 0.001 |
| **cg21879725** | **12.0** | **0.002** | **0.001** | **18.0** | **1.3 x 10^-4^** | **0.003** |

**Supp. Table 3:** Basic- and fully-adjusted linear mixed models with methylation level as the outcome variable and age as the predictor variable. Bold font denotes significant values after correcting for multiple comparisons (p < 0.0038). The CpGs are classified into region 1 (orange), region 2 (green), or region 3 (blue).

|  | **Basic-adjusted model** | | | **Fully-adjusted model** | | |
| --- | --- | --- | --- | --- | --- | --- |
| **CpG** | **Estimate** | **SE** | **p** | **Estimate** | **SE** | **p** |
| **cg20051876** | **-0.016** | **9.7 x 10^-4^** | **<2.0 x 10^-16^** | **-0.011** | **0.0014** | **<2.0 x 10^-16^** |
| **cg14123992** | **-0.017** | **0.0011** | **<2.0 x 10^-16^** | **-0.018** | **0.0015** | **<2.0 x 10^-16^** |
| **cg04406254** | **-0.0034** | **10.0 x 10^-4^** | **5.4 x 10^-4^** | -0.0025 | 0.0014 | 0.07 |
| cg26190885 | 9.8 x 10^-5^ | 0.0013 | 0.94 | -0.0011 | 0.0019 | 0.55 |
| **cg12049787** | **0.0037** | **0.0012** | **0.002** | 0.0031 | 0.0017 | 0.06 |
| **cg08955609** | **0.0054** | **0.0014** | **1.3 x 10^-4^** | 0.0048 | 0.0019 | 0.01 |
| **cg18768621** | **0.0091** | **0.0012** | **1.4 x 10^-13^** | **0.0092** | **0.0017** | **6.9 x 10^-8^** |
| cg19514613 | 0.0028 | 0.0013 | 0.03 | 0.0039 | 0.0018 | 0.03 |
| cg06750524 | 0.0016 | 0.0015 | 0.45 | 4.4 x 10^-4^ | 0.0021 | 0.84 |
| cg16471933 | -0.0031 | 0.0011 | 0.004 | -0.0027 | 0.0015 | 0.08 |
| **cg05501958** | **-0.0038** | **0.0011** | **3.7 x 10^-4^** | -0.0042 | 0.0015 | 0.0039 |
| cg18799241 | 5.6 x 10^-5^ | 8.9 x 10^-4^ | 0.95 | 6.1 x 10^-4^ | 0.0013 | 0.63 |
| cg21879725 | -0.0016 | 0.0010 | 0.14 | -0.0017 | 0.0015 | 0.26 |

**Supp. Table 4**: Breusch-Pagan test for heteroscedasticity for the basic- and fully-adjusted model. Bold font denotes significant values after correcting for multiple comparisons (p < 0.0038). The CpGs are classified into region 1 (orange), region 2 (green), or region 3 (blue).

|  | **Basic-adjusted model** | | **Fully-adjusted model** | |
| --- | --- | --- | --- | --- |
| **CpG** | **BP** | **p** | **BP** | **p** |
| **cg20051876** | **120.4** | **9.3 x 10^-11^** | **77.3** | **7.4 x 10^-4^** |
| **cg14123992** | **142.6** | **2.6 x 10^-14^** | **87.5** | **4.8 x 10^-5^** |
| **cg04406254** | **225.2** | **2.2 x 10^-16^** | **140.6** | **1.5 x 10^-12^** |
| **cg26190885** | **497.5** | **<2.2 x 10^-16^** | **218.5** | **<2.2 x 10^-16^** |
| **cg12049787** | **99.5** | **1.2 x 10^-7^** | **98.6** | **1.9 x 10^-6^** |
| **cg08955609** | **968.6** | **<2.2 x 10^-16^** | **125.1** | **<2.2 x 10^-16^** |
| **cg18768621** | **931.4** | **<2.2 x 10^-16^** | **302.7** | **<2.2 x 10^-16^** |
| **cg19514613** | **736.7** | **<2.2 x 10^-16^** | **185.2** | **<2.2 x 10^-16^** |
| **cg06750524** | **301.7** | **<2.2 x 10^-16^** | **155.1** | **7.2 x 10^-15^** |
| **cg16471933** | **535.3** | **<2.2 x 10^-16^** | **177.1** | **<2.2 x 10^-16^** |
| **cg05501958** | **491.3** | **<2.2 x 10^-16^** | **258.1** | **<2.2 x 10^-16^** |
| **cg18799241** | **295.4** | **<2.2 x 10^-16^** | **142.0** | **8.9 x 10^-13^** |
| **cg21879725** | **378.1** | **<2.2 x 10^-16^** | **241.1** | **<2.2 x 10^-16^** |

**Supp. Table 5**: Basic- and fully-adjusted linear mixed models with methylation level as the outcome variable and cognitive-g as the predictor variable. Bold font denotes significant values after correcting for multiple comparisons (p < 0.0038). The CpGs are classified into region 1 (orange), region 2 (green), or region 3 (blue).

|  | **Basic-adjusted model** | | | **Fully-adjusted model** | | |
| --- | --- | --- | --- | --- | --- | --- |
| **CpG** | **Estimate** | **SE** | **p** | **Estimate** | **SE** | **p** |
| cg20051876 | 0.0029 | 0.0084 | 0.73 | -0.0045 | 0.013 | 0.73 |
| cg14123992 | -0.0083 | 0.0092 | 0.37 | -0.0012 | 0.014 | 0.93 |
| cg04406254 | 0.0082 | 0.0084 | 0.33 | 0.0038 | 0.013 | 0.77 |
| cg26190885 | 0.012 | 0.011 | 0.33 | -0.023 | 0.018 | 0.19 |
| cg12049787 | 0.0012 | 0.010 | 0.91 | 0.016 | 0.016 | 0.32 |
| cg08955609 | 0.013 | 0.012 | 0.29 | 0.025 | 0.018 | 0.18 |
| cg18768621 | -0.015 | 0.011 | 0.15 | -0.0062 | 0.016 | 0.70 |
| cg19514613 | -0.0082 | 0.011 | 0.45 | -0.0062 | 0.017 | 0.72 |
| **cg06750524** | **0.044** | **0.013** | **6.2 x 10^-4^** | 0.039 | 0.020 | 0.05 |
| cg16471933 | 0.017 | 0.0093 | 0.07 | 0.012 | 0.015 | 0.43 |
| cg05501958 | 0.009 | 0.0090 | 0.92 | -0.0029 | 0.014 | 0.84 |
| cg18799241 | 0.011 | 0.0076 | 0.17 | 0.0042 | 0.012 | 0.72 |
| cg21879725 | 0.015 | 0.0088 | 0.09 | 0.030 | 0.014 | 0.035 |

**Supp. Table 6:** (also next page): Basic- and fully-adjusted linear mixed models with methylation level as the outcome variable and individual cognitive tests as predictor variables. Bold font denotes significant values after correcting for multiple comparisons (p < 9.6 x 10^-4^). The CpGs are classified into region 1 (orange), region 2 (green), or region 3 (blue).

| **Basic-adjusted model** | | | | | | | | |
| --- | --- | --- | --- | --- | --- | --- | --- | --- |
|  | **Dig.** | | **Verbal** | | **M.-H.** | | **Log.** | |
| **CpG** | **estimate (SE)** | **p** | **estimate (SE)** | **p** | **estimate (SE)** | **p** | **estimate (SE)** | **p** |
| **cg20051876** | **0.045 (0.0091)** | **5.5 x 10^-7^** | -0.012 (0.0093) | 0.20 | **-0.040 (0.0094)** | **1.9 x 10^-5^** | 0.018 (0.0090) | 0.050 |
| **cg14123992** | **0.047 (0.0099)** | **1.9 x 10^-6^** | -0.0017 (0.010) | 0.87 | **-0.076 (0.010)** | **1.1 x 10^-13^** | 0.027 (0.0098) | 0.006 |
| cg04406254 | 0.020 (0.0092) | 0.03 | -0.0088 (0.0094) | 0.35 | -0.0092 (0.0095) | 0.33 | 0.014 (0.0091) | 0.13 |
| cg26190885 | 0.0097 (0.012) | 0.44 | 0.0075 (0.013) | 0.56 | 0.013 (0.013) | 0.30 | -0.016 (0.012) | 0.19 |
| cg12049787 | -0.025 (0.011) | 0.02 | -0.0029 (0.011) | 0.80 | 0.015 (0.011) | 0.18 | 0.014 (0.011) | 0.21 |
| cg08955609 | 0.0032 (0.013) | 0.81 | 0.0049 (0.013) | 0.71 | 0.032 (0.013) | 0.02 | -0.023 (0.013) | 0.07 |
| **cg18768621** | **-0.039 (0.011)** | **5.4 x 10^-4^** | 0.0073 (0.012) | 0.53 | 0.011 (0.012) | 0.35 | -0.0059 (0.011) | 0.60 |
| cg19514613 | 0.0012 (0.012) | 0.92 | -0.013 (0.012) | 0.28 | 0.016 (0.012) | 0.20 | -0.017 (0.012) | 0.15 |
| cg06750524 | 0.0096 (0.014) | 0.49 | 0.022 (0.014) | 0.14 | 0.033 (0.015) | 0.02 | -3.9 x 10^-4^ (0.014) | 0.98 |
| cg16471933 | 0.011 (0.010) | 0.26 | 0.0095 (0.010) | 0.36 | 0.0087 (0.010) | 0.40 | -0.0043 (0.010) | 0.67 |
| cg05501958 | 0.024 (0.0098) | 0.01 | -0.015 (0.010) | 0.13 | -0.056 (0.010) | 0.58 | -0.0018 (0.0097) | 0.86 |
| cg18799241 | -0.0030 (0.0082) | 0.72 | 0.012 (0.0084) | 0.20 | -0.0053 (0.0085) | 0.53 | 0.013 (0.0081) | 0.10 |
| cg21879725 | 0.0086 (0.0096) | 0.37 | 0.011 (0.0099) | 0.25 | -0.0087 (0.0095) | 0.38 | 0.013 (0.0095) | 0.18 |

| **Fully-adjusted model** | | | | | | | | |
| --- | --- | --- | --- | --- | --- | --- | --- | --- |
|  | **Dig.** | | **Verbal** | | **M.-H.** | | **Log.** | |
| **CpG** | **estimate (SE)** | **p** | **estimate (SE)** | **p** | **estimate (SE)** | **p** | **estimate (SE)** | **p** |
| cg20051876 | 0.0093 (0.013) | 0.49 | -0.0086 (0.013) | 0.49 | 0.011 (0.014) | 0.45 | 8.5 x 10^-4^ (0.012) | 0.94 |
| cg14123992 | -0.017 (0.014) | 0.23 | 0.021 (0.013) | 0.12 | -0.0064 (0.015) | 0.68 | 0.0026 (0.013) | 0.85 |
| cg04406254 | 0.018 (0.014) | 0.18 | -0.010 (0.013) | 0.43 | -0.0050 (0.015) | 0.73 | 0.0037 (0.012) | 0.77 |
| cg26190885 | 0.029 (0.018) | 0.11 | -0.010 (0.017) | 0.55 | -0.037 (0.020) | 0.06 | -0.013  (0.017) | 0.44 |
| cg12049787 | -0.0095 (0.016) | 0.56 | 0.0056 (0.015) | 0.72 | -0.0061 (0.018) | 0.73 | 0.027  (0.015) | 0.07 |
| cg08955609 | 0.033 (0.019) | 0.08 | -0.0017 (0.018) | 0.93 | 0.014 (0.020) | 0.50 | -1.7 x 10^-4^ (0.017) | 0.99 |
| cg18768621 | -0.0051 (0.017) | 0.76 | 0.0033 (0.016) | 0.83 | 1.3 x 10^-4^ (0.018) | 0.99 | 0.0022 (0.015) | 0.88 |
| cg19514613 | -0.0018 (0.018) | 0.92 | -0.0058 (0.017) | 0.72 | 0.011 (0.019) | 0.55 | -0.0063 (0.016) | 0.70 |
| cg06750524 | 0.015 (0.021) | 0.45 | 0.038 (0.020) | 0.06 | -0.0067 (0.023) | 0.77 | 0.0057 (0.019) | 0.77 |
| cg16471933 | -0.0034 (0.015) | 0.82 | -9.2 x 10^-4^ (0.014) | 0.95 | 0.019 (0.016) | 0.25 | 0.0014 (0.014) | 0.92 |
| cg05501958 | 0.020 (0.014) | 0.16 | -0.020 (0.014) | 0.13 | 0.012 (0.016) | 0.45 | -0.0037 (0.013) | 0.78 |
| cg18799241 | -0.0089 (0.012) | 0.47 | -0.0024 (0.012) | 0.84 | 0.0060 (0.013) | 0.65 | 0.011  (0.011) | 0.32 |
| cg21879725 | -0.020 (0.015) | 0.17 | 0.030 (0.014) | 0.03 | 0.0085 (0.016) | 0.59 | 0.018  (0.013) | 0.19 |

**Supp. Table 7**: Basic- and fully-adjusted linear mixed models with methylation level as the outcome variable and AD-risk as the predictor variable. Bold font denotes significant values after correcting for multiple comparisons (p < 0.0038). The CpGs are classified into region 1 (orange), region 2 (green), or region 3 (blue).

|  | **Basic-adjusted model** | | | **Fully-adjusted model** | | | **Fully-adjusted model + genotype** | | |
| --- | --- | --- | --- | --- | --- | --- | --- | --- | --- |
| **CpG** | **Estimate** | **(SE)** | **p** | **Estimate** | **(SE)** | **p** | **Estimate** | **(SE)** | **p** |
| cg20051876 | -0.0075 | (0.0083) | 0.36 | -0.0012 | 0.011 | 0.92 | -0.0033 | 0.011 | 0.77 |
| cg14123992 | -0.0091 | (0.0091) | 0.32 | 0.016 | 0.012 | 0.19 | 0.013 | 0.012 | 0.29 |
| cg04406254 | -0.0010 | (0.0083) | 0.90 | -0.0013 | 0.011 | 0.91 | -0.0089 | 0.011 | 0.43 |
| cg26190885 | -9.2 x 10^-4^ | (0.011) | 0.94 | 0.0017 | 0.015 | 0.91 | -9.0 x 10^-5^ | 0.015 | 1.0 |
| cg12049787 | 0.014 | (0.010) | 0.16 | 0.0029 | 0.013 | 0.83 | 0.0043 | 0.014 | 0.76 |
| cg08955609 | 0.0041 | (0.012) | 0.73 | 0.0051 | 0.015 | 0.74 | 0.0038 | 0.014 | 0.79 |
| cg18768621 | 0.0037 | (0.010) | 0.72 | 0.0052 | 0.014 | 0.70 | 0.0038 | 0.014 | 0.79 |
| cg19514613 | 0.0079 | (0.011) | 0.47 | 0.021 | 0.015 | 0.14 | 0.016 | 0.015 | 0.27 |
| cg06750524 | 0.0072 | (0.033) | 0.57 | 0.0018 | 0.017 | 0.92 | -0.023 | 0.017 | 0.19 |
| cg16471933 | 0.024 | (0.0092) | 0.009 | 0.035 | 0.012 | 0.004 | 0.025 | 0.013 | 0.04 |
| cg05501958 | 0.013 | (0.0089) | 0.14 | 0.018 | 0.012 | 0.14 | 0.018 | 0.012 | 0.13 |
| cg18799241 | 0.0013 | (0.0075) | 0.87 | -0.0033 | 0.010 | 0.75 | -3.3 X 10^-4^ | 0.010 | 0.97 |
| cg21879725 | 0.011 | (0.0088) | 0.20 | 0.0030 | 0.012 | 0.80 | -0.0027 | 0.012 | 0.83 |

**Supp. Table 8**: Basic- and fully-adjusted linear mixed models with methylation level as the outcome variable and CVD-risk as the predictor variable. Bold font denotes significant values after correcting for multiple comparisons (p < 0.0038). The CpGs are classified into region 1 (orange), region 2 (green), or region 3 (blue).

|  | **Basic-adjusted model** | | | **Fully-adjusted model** | | |
| --- | --- | --- | --- | --- | --- | --- |
|  | **Estimate** | **SE** | **p** | **Estimate** | **SE** | **p** |
| cg20051876 | -0.021 | 0.083 | 0.010 | -0.0015 | 0.012 | 0.89 |
| cg14123992 | -0.013 | 0.0091 | 0.15 | 0.0066 | 0.012 | 0.60 |
| cg04406254 | -0.0037 | 0.0083 | 0.66 | -0.014 | 0.012 | 0.24 |
| cg26190885 | -0.029 | 0.011 | 0.010 | -0.020 | 0.016 | 0.21 |
| cg12049787 | 0.012 | 0.0099 | 0.24 | -0.0064 | 0.014 | 0.65 |
| cg08955609 | -0.022 | 0.012 | 0.042 | -0.012 | 0.016 | 0.46 |
| cg18768621 | -0.0021 | 0.010 | 0.84 | 0.0034 | 0.014 | 0.81 |
| cg19514613 | 0.0034 | 0.011 | 0.75 | 0.013 | 0.015 | 0.40 |
| cg06750524 | 0.0032 | 0.013 | 0.80 | -0.0022 | 0.018 | 0.90 |
| cg16471933 | -0.0044 | 0.0091 | 0.63 | -0.0026 | 0.013 | 0.84 |
| cg05501958 | -0.0022 | 0.0089 | 0.81 | 0.016 | 0.012 | 0.21 |
| cg18799241 | 0.0024 | 0.0074 | 0.74 | 9.5 x 10^-5^ | 0.011 | 0.99 |
| cg21879725 | 0.021 | 0.0087 | 0.017 | 0.011 | 0.013 | 0.38 |

**Supp. Table 9**: Basic- and fully-adjusted linear mixed models and the fully-adjusted linear mixed model with the addition of genotype as a covariate, with methylation level as the outcome variable and the quotient of total cholesterol and HDL-cholesterol as the predictor variable. Bold font denotes significant values after correcting for multiple comparisons (p < 0.0038). The CpGs are classified into region 1 (orange), region 2 (green), or region 3 (blue).

|  | **Basic-adjusted model** | | | **Fully-adjusted model** | | | **Fully-adjusted mode + genotype** | | |
| --- | --- | --- | --- | --- | --- | --- | --- | --- | --- |
| **CpG** | **Estimate** | **SE** | **p** | **Estimate** | **SE** | **p** | **Estimate** | **SE** | **p** |
| cg20051876 | -0.020 | 0.0083 | 0.01 | 0.013 | 0.013 | 0.32 | 0.0067 | 0.013 | 0.62 |
| cg14123992 | -0.0058 | 0.0092 | 0.53 | 0.020 | 0.014 | 0.16 | 0.0067 | 0.014 | 0.64 |
| cg04406254 | 0.0041 | 0.0084 | 0.63 | 0.010 | 0.013 | 0.45 | 0.0021 | 0.014 | 0.87 |
| cg26190885 | -0.017 | 0.011 | 0.14 | -0.023 | 0.018 | 0.20 | -0.023 | 0.018 | 0.20 |
| cg12049787 | 0.014 | 0.010 | 0.18 | -0.014 | 0.016 | 0.38 | -0.017 | 0.017 | 0.30 |
| **cg08955609** | **-0.046** | **0.012** | **1.2 x 10^-4^** | **-0.064** | **0.019** | **5.3 x 10^-4^** | **-0.065** | **0.019** | **6.7 x 10^-4^** |
| **cg18768621** | **-0.048** | **0.010** | **4.8 x 10^-6^** | **-0.056** | **0.016** | **6.7 x 10^-4^** | **-0.059** | **0.017** | **4.3 x 10^-4^** |
| **cg19514613** | **-0.050** | **0.011** | **6.1 x 10^-6^** | -0.040 | 0.017 | 0.02 | -0.051 | 0.018 | 0.0042 |
| cg06750524 | 0.0047 | 0.013 | 0.72 | 0.0050 | 0.021 | 0.81 | -0.025 | 0.021 | 0.22 |
| **cg16471933** | **0.048** | **0.0092** | **2.4 x 10^-7^** | **0.059** | **0.015** | **6.7 x 10^-5^** | 0.036 | 0.015 | 0.02 |
| cg05501958 | 9.0 x 10^-4^ | 0.0089 | 0.92 | 0.0090 | 0.014 | 0.52 | 0.011 | 0.014 | 0.45 |
| cg18799241 | -0.0023 | 0.0075 | 0.76 | -6.1 x 10^-4^ | 0.012 | 0.96 | 0.0033 | 0.012 | 0.79 |
| cg21879725 | -0.0016 | 0.0089 | 0.85 | -0.0054 | 0.014 | 0.71 | -0.015 | 0.017 | 0.30 |
